# MosaicBase: A Knowledgebase of Postzygotic Mosaic Variants in Noncancer Diseases and Asymptomatic Human Individuals

**DOI:** 10.1101/2020.03.04.976803

**Authors:** Xiaoxu Yang, Changhong Yang, Xianing Zheng, Luoxing Xiong, Yutian Tao, Meng Wang, Adam Yongxin Ye, Qixi Wu, Yanmei Dou, Junyu Luo, Liping Wei, August Yue Huang

## Abstract

Mosaic variants resulting from postzygotic mutations are prevalent in the human genome and play important roles in human diseases. However, except for cancer-related variant collections, there are no collections of mosaic variants in noncancer diseases and asymptomatic individuals. Here, we present MosaicBase (http://mosaicbase.cbi.pku.edu.cn/ or http://49.4.21.8:8000/), a comprehensive database that includes 6,698 mosaic variants related to 269 noncancer diseases and 27,991 mosaic variants identified in 422 asymptomatic individuals. The genomic and phenotypic information for each variant was manually extracted and curated from 383 publications. MosaicBase supports the query of variants with Online Mendelian Inheritance in Man (OMIM) entries, genomic coordinates, gene symbols, or Entrez IDs. We also provide an integrated genome browser for users to easily access mosaic variants and their related annotations within any genomic region. By analyzing the variants collected in MosaicBase, we found that mosaic variants that directly contribute to disease phenotype showed features distinct from those of variants in individuals with a mild or no phenotype in terms of their genomic distribution, mutation signatures, and fraction of mutant cells. MosaicBase will not only assist clinicians in genetic counseling and diagnosis but also provide a useful resource to understand the genomic baseline of postzygotic mutations in the general human population.

## Introduction

Genomic mosaicism results from postzygotic mutations arising during embryonic development, tissue self-renewal [1], aging processes [2], or exposure to other DNA-damaging circumstances [3]. Unlike *de novo* or inherited germline variants that affect every cell in the carrier individual [4], postzygotic mosaic variants only affect a portion of cells or cell populations, and their mutant allelic fractions (MAFs) should be 50% [5]. If a postzygotic mutation affects germ cells [6], the mutant allele may theoretically be transmitted to offspring, which is the major source of genetic variations in the human population [7].

Postzygotic mosaic variants have previously been demonstrated to be directly responsible for the etiology of cancer [8, 9] and an increasing number of other Mendelian or complex diseases, including epilepsy-related neurodevelopment disorders [10], Costello syndrome [11], autism spectrum disorders [12, 13], and intellectual disability [14]. On the other hand, pathogenic genetic variants inherited from detectable parental mosaicism have been demonstrated to be an important source of monogenic genetic disorders, including Noonan syndrome [15], Marfan syndrome [16], Dravet syndrome [17], and complex disorders, including autism [18] and intellectual disability [19]. The MAF of a mosaic variant has been reported to be directly related to the carrier’s phenotype [20, 21] and to be associated with the recurrence risk in children [5].

With the rapid advances in next-generation sequencing (NGS) technologies, tens of thousands of postzygotic mosaic single-nucleotide variants (SNVs) and insertions/deletions (indels) have been identified and validated in the genomes of human individuals [3, 22, 23]. However, except for cancer-related variants that have been collected by databases such as the Catalogue of Somatic Mutations in Cancer (COSMIC) [24] and SomamiR (somatic mutations impacting microRNA function in cancer) [25], there is no integrated database focusing on mosaic variants in noncancer diseases and asymptomatic individuals.

Here, we present MosaicBase (http://mosaicbase.cbi.pku.edu.cn/ or http://49.4.21.8:8000/); to our knowledge, MosaicBase is the first knowledgebase of mosaic SNVs and indels identified in patients with noncancer diseases and their parents as well as asymptomatic individuals. MosaicBase currently contains 34,689 validated mosaic variants that have been manually curated from 383 publications. MosaicBase has further integrated comprehensive genomic and phenotypic information about each variant and its carrier. It provides multi-scale information about disease-related mosaic variants for genetic counseling and molecular diagnosis as well as the genomic background of mosaic variants in general populations.

## Database implementation

### The framework of MosaicBase

An overview of the framework of MosaicBase is shown in Figure 1. MosaicBase consists of two logical parts: the database and server as the backend and the user interface as the frontend. Structured data based on three relational tables were established in the backend of MosaicBase. The storage and maintenance of the database were implemented with SQLite v3. The frontend of MosaicBase provides a user-friendly interface written in PHP, JavaScript, HTML and CSS, with Django applications.

**Figure 1:**
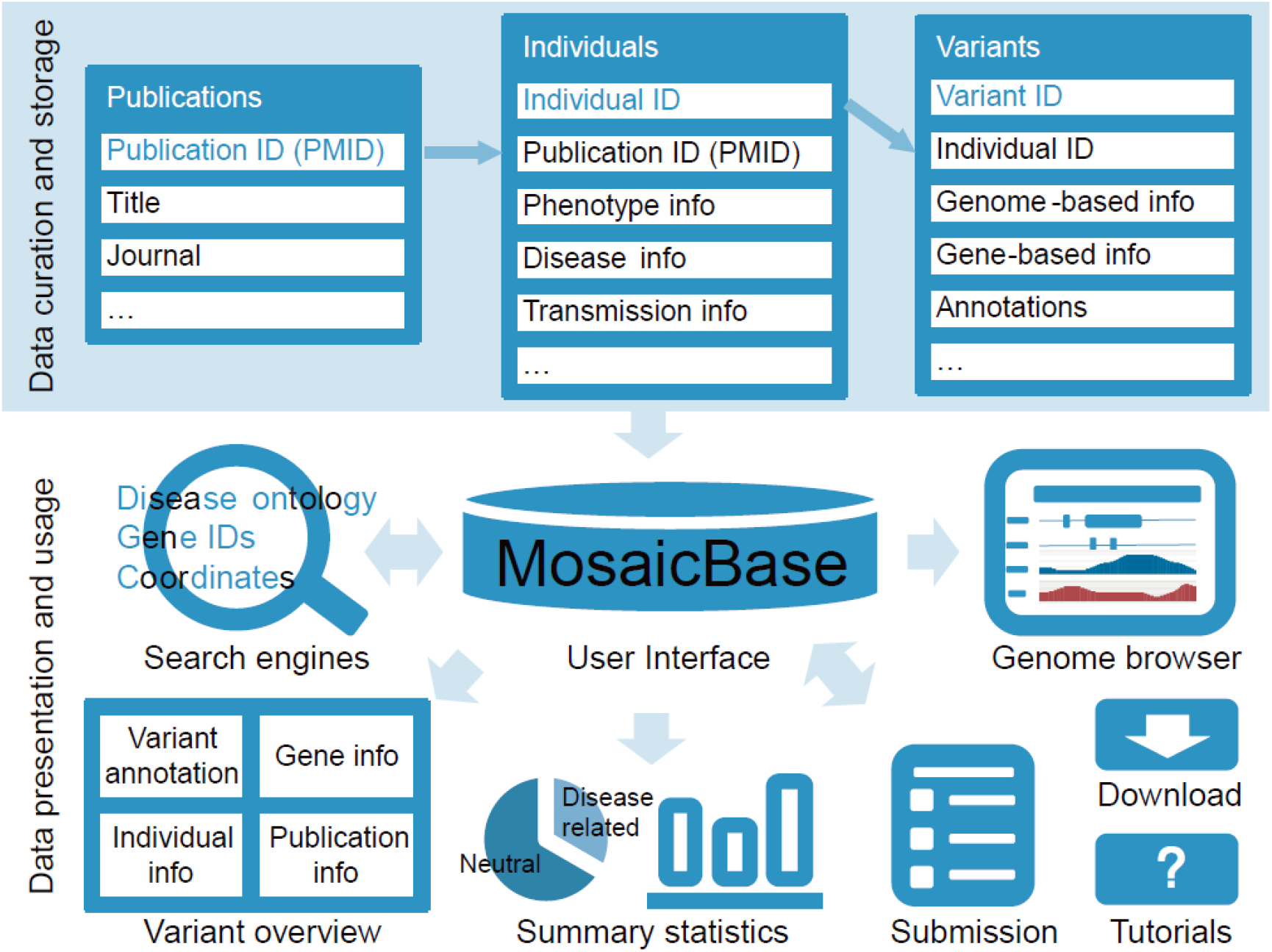
Overview of the data collection, storage, and visualization of MosaicBase.

MosaicBase incorporates two different search modes to help the user browse the database. The information for each mosaic variant has been summarized from the publication and individual levels to the gene and variant levels. A built-in genome browser is provided to visualize variants. A statistical summary and detailed tutorials for MosaicBase are available on the main page. MosaicBase further provides an online submission system to encourage the community to contribute to the database.

### Data collection, processing, and annotation

We queried against the PubMed database using keywords including “mosaic”, “mosaicism”, “post-zygotic”, “somatic”, “sequencing” (see the full query string in Supplemental Text), and excluded publications about cancer-related mosaic mutations or studies on non-human organisms by examining the titles and abstracts. For more than 1,000 search results, we scrutinized the main text as well as supplemental information to confirm the relevance of each publication. After this process, 383 journal research articles about mosaic SNVs and indels in noncancer individuals that were published between Jan 1989 and May 2018 were collected into MosaicBase. For each article, data fields for the publication, individual, and variation information were extracted and saved into three tables in the backend (Figure 1). For studies involving single-cell technologies, only the validated or high-confidence postzygotic mosaic SNVs were collected. For the table of variation information, we further integrated the genomic annotations generated by ANNOVAR [26], including population allele frequency from dbSNP (version 137) [27] and gnomAD (genome; version 2.0.1) [28], risk scores such as CADD scores (version 1.30) [29] and Eigen scores [30], functional predictions by FATHMM [31], SIFT [32], iFish2 [33], DeFine [34], conservation prediction by GERP++ [35] and PhyloP [36], and annotations in COSMIC [37]. A detailed description of different fields and data types required in each field is listed in Supp. Tables S1, S2, and S3. The transcript-based variation information was confirmed using Mutalyzer following the suggestions from the Human Genome Variation Society (HGVS) [38]. Genomic coordinates were provided according to the human reference genome UCSC hg19/GRCh37 as well as hg38/GRCh38.

### Statistical analysis and visualization of mosaic variants

The mutation signature analysis has been widely used in cancer studies to elaborate the etiology of somatic mosaic variants, by decomposing the matrix of tri-nucleotide context into cancer-related signatures. In this study, the signature of noncancer mosaic variants was analyzed by Mutalisk [39], and the maximum likelihood estimation of proportions for each mutation signature was performed based on a greedy algorithm. For each variant group, we further tested whether its genomic density within each 1 Mb interval was correlated with the GC content, DNase I hypersensitive regions, replication timing, and histone modification profiles measured in the GM12878 cell line [39]. A genome browser based on the Dalliance platform [40] was implemented to interactively visualize the mosaic variants. Circos [41] was utilized to show the genomic distribution of mosaic variants.

## Web interface

### User interface and functions

We incorporated two search modes in MosaicBase. The basic search mode provided on the main page recognizes search terms based on the name of diseases, the range of genomic coordinates, gene symbols, or Entrez Gene IDs (Figure 2A), in which the search engine is comparable with space-delimited multiple search terms. The result page of the basic search mode displays variant summary information according to the categories of search terms, and search results can then be downloaded as an xls format table. We also introduced an ontology-based search mode as an advanced option in MosaicBase; with this mode, users can browse the mosaic variants related to a specific disease or disease category according to the Disease Ontology [42]. A brief summary of the description of the disease or disease category is provided along with a summary table of all the related mosaic variants collected in MosaicBase (Figure 2B).

**Figure 2:**
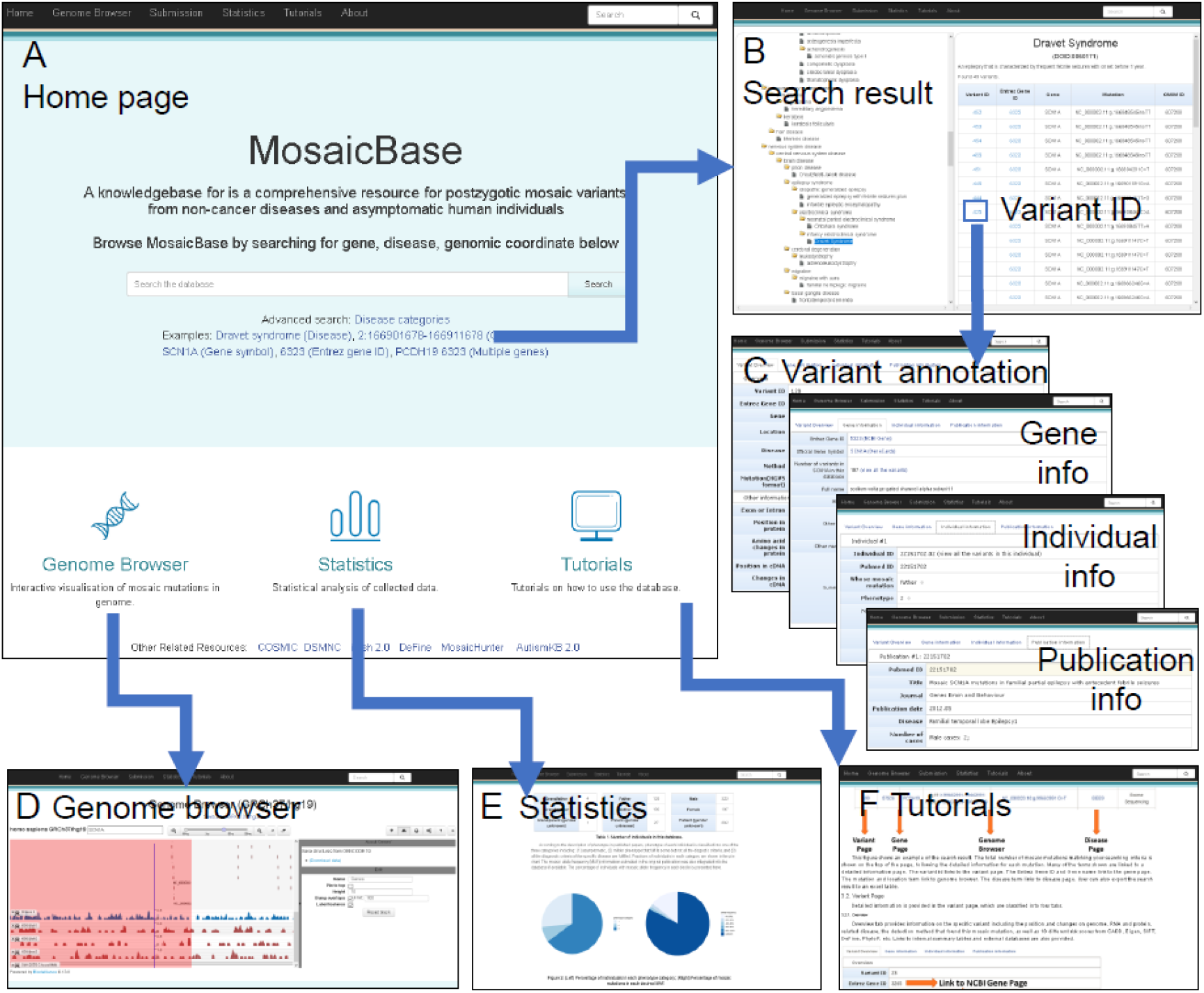
Screenshots of MosaicBase. **A.** The main page provides the search modes and multiple links to different utilities of the database. **B.** Disease-ontology-based advanced search page and an example of a result table. **C.** The variant pages from the basic search results; this page provides information about each variant and its corresponding gene, individual and publication annotation, the individuals carrying the same variant, and the publication describing the variant. **D.** Summary statistics of the publications, mutational spectrum, and individuals collected in MosaicBase. **E.** Integrated genome browser to visualize mosaic variants with genetic and epigenetic annotations. **F.** Detailed tutorials for the introduction, data presentation, and usage of MosaicBase.

Detailed information about each mosaic variant was summarized in four different panels in MosaicBase: the overview panel, the gene information panel, the individual information panel, and the publication information panel (Figure 2C). In the overview panel, we provided the genomic information as well as the methodologies for the identification and validation of the variant. In the gene information panel, we annotated the Entrez Gene ID, official gene symbol and alternative names, number of reported mosaic variants in this gene, Vega ID, OMIM ID, HGNC ID, Ensembl ID, and a brief summary of the gene. In the gene information panel, we summarized all the collected mosaic variants in the same gene and provided various resources for gene annotation from external databases, including Entrez, Vega, OMIM, HGNC, and Ensembl IDs. In the individual information panel, we classified the phenotypes of the individual carrying the mosaic variant and displayed the information according to the original descriptions in the publication. The severity of phenotype collected in MosaicBase was defined as “1” if the carrier was asymptomatic, “2” if the carrier had a mild phenotype but did not fulfill all the diagnostic criteria for a specific disease or characterized syndrome, and “3” if the carrier fulfilled all the clinical diagnostic criteria for a specific disease. In the publication information panel, we summarized the title, journal, sample, and additional information about the publication that reported the mosaic variant.

MosaicBase integrated a build-in genome browser to provide convenient interactive data visualization for the mosaic variants (Figure 2D). In addition to the default tracks about genetic and epigenetic annotations, such as DNase I hypersensitive sites and H3K4me predictions, MosaicBase also allows the user to import customized tracks from URLs, UCSC-style track hubs, or uploaded files in a UCSC-style genome browser track format. The URLs for tracks of Ensembl Gene and MeDIP-seq data are provided as examples, and a help page providing detailed guidance is also available by clicking the question mark in the top-left panel of the genome browser. MosaicBase further provided users with an application that can generate publication-quality SVG files from the control panel of the genome browser.

MosaicBase included a “Statistics” page to show a summary of all the collected mosaic variants (Figure 2E) and a “Tutorials” page (Figure 2F) with detailed introductions about the database and its search modes, data presentation, and genome browser. We also implemented an online submission system that allows users to submit mosaic variants from newly published or uncollected publications. Such variants will be manually examined by our team and integrated into MosaicBase with scheduled updates.

### Statistical analysis of noncancer mosaic variants

MosaicBase currently includes 383 journal research articles, letters, and clinical genetic reports about noncancer postzygotic mosaic variants that were published from 1989 to 2018 (Figure 3A), with an accelerated accumulation of mosaic-related publications boosted by the recent advances in NGS technologies. After manually extracting the mosaic variants reported in each publication, we thoroughly compiled 34,689 mosaic variants from 2,202 noncancer individuals, including 6,698 disease-related variants from 3,638 genes related to 269 noncancer diseases as well as 27,991 apparently neutral variants identified from 442 asymptomatic individuals (Figure 3B and Supp. Table S4). Specifically, two types of disease-related mosaic variants were collected in MosaicBase: 1) 6,207 mosaic variants that had directly contributed to the disease phenotype in 1,402 patients (323 men and 197 women; 882 sex unknown from the original publication) and 2) 491 mosaic variants identified from 358 parents or grandparents (137 men and 193 women; 28 sex unknown from the original publication) of the probands who had transmitted the mosaic allele to their offspring for a heterozygous genotype that led to disease phenotypes (Figure 3B). The collected mosaic variants were classified into three groups according to the origin of the variants described in the original publications: variants from asymptomatic individuals were termed the “asym” group; variants from patients fulfilling the full diagnostic criteria of a specific disease were termed the “patient” group; variants from parents/grandparents of the patients were termed the “parent” group. As shown in Figure 3C, mosaic variants were generally distributed across all the autosomes and X chromosomes. Parental mosaic variants were clustered in the *SCN1A* gene on chromosome 2, which resulted from the well-studied parental mosaic cases for Dravet syndrome. The underrepresentation of mosaic variants in the Y chromosome might be explained by its low gene density and the technical challenge of detecting mosaic variants in haplotype chromosomes.

**Figure 3:**
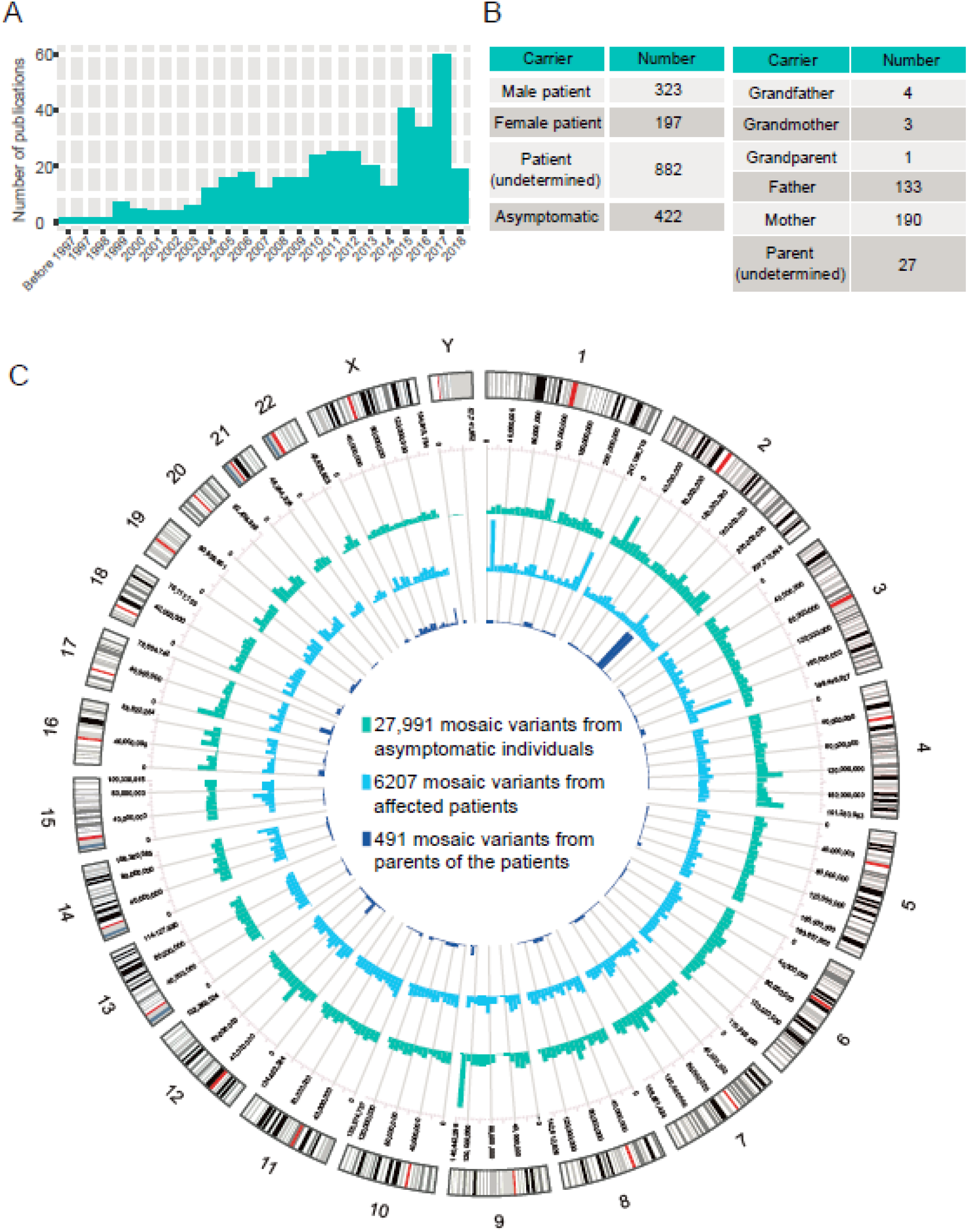
Statistics about the publication, individual, and variant data collected in MosaicBase. **A.** Number of mosaic-related publications from 1989 to 2018. **B.** Summary of different categories of mosaic carriers. **C.** Circos plot of mosaic variants. Histograms show the number of mosaic variants for each 1 Mb genomic window. Chromosomal bands are illustrated in the outer circle with centromeres in red.

To study whether mosaic variants from different groups of individuals have distinct genomic characteristics, we calculated their correlation with various genomic regulation features, including GC content, DNase I hypersensitive positions, and epigenetic modifications. Because the vast majority of mosaic variants had been identified from peripheral blood or saliva samples, genomic regulation patterns of GM12878, a lymphoblastoid-derived cell line, were used in the subsequent analysis. Common germline variants annotated in dbSNP 137 with allele frequency higher than 10% (“dbSNP” group) were served as a control. According to the Pearson correlation coefficients between the signal intensities of genomic features and the density of variants with a window size of 1MB across the genome [43], we found that the mosaic variants that directly contribute to the disease phenotype (“patient” group) are more positively correlated with such genomic features than the mosaic variants of the other groups (Figure 4A).

**Figure 4:**
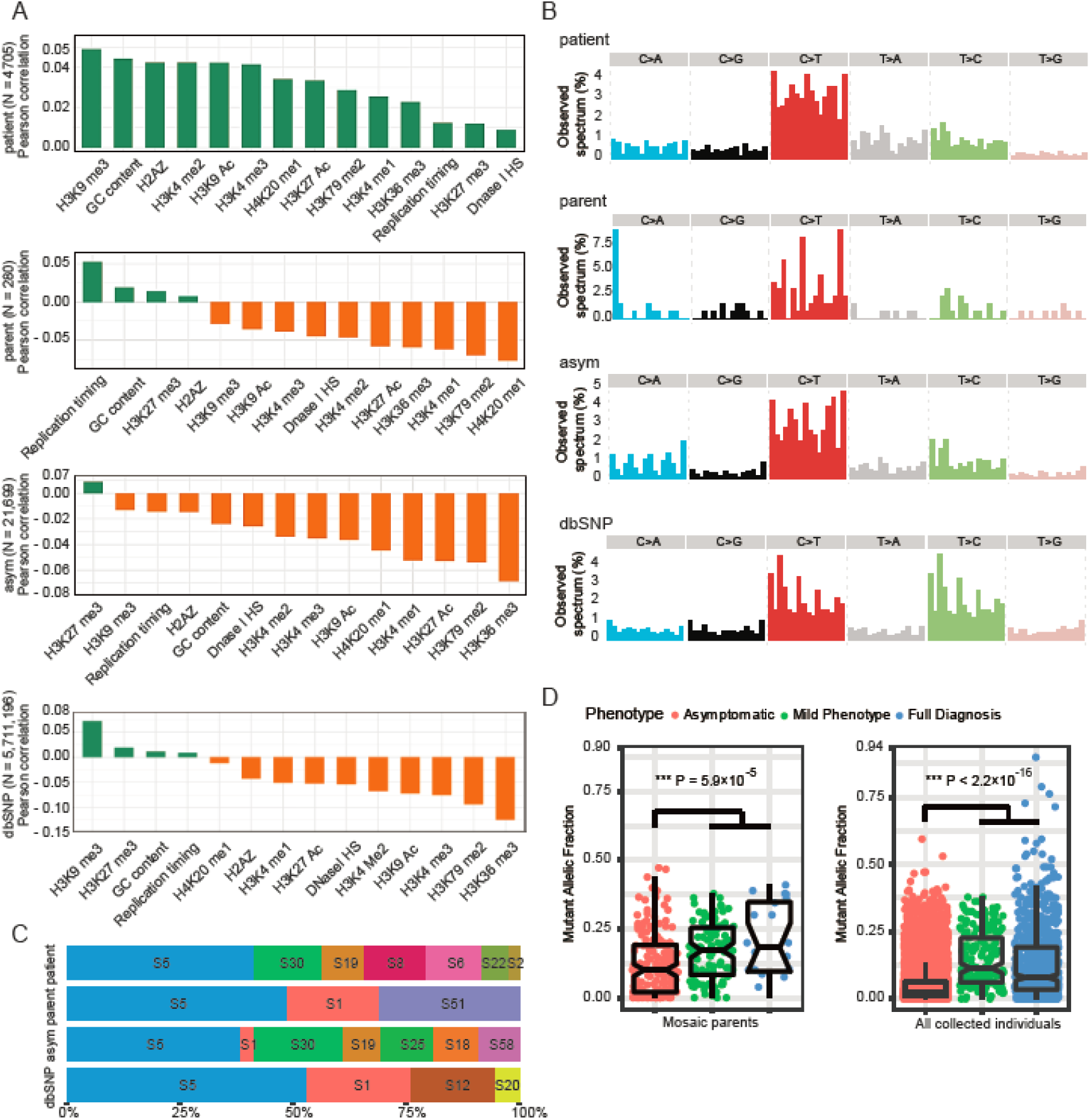
Genomic features of mosaic variants collected in MosaicBase. **A.** Correlation of the density of mosaic variants and various genomic regulation features. **B.** Tri-nucleotide genomic context of mosaic variants. **C.** Proportion of cancer signatures for mosaic variants. **D.** Mutant allele fraction of mosaic variants in mosaic parents only (P = 5.9×10^−5^ by a Mann–Whitney U test with continuity correction, left) and in all individuals (P < 2.2×10^−16^ by a Mann–Whitney U test, right). Common germline variants with population allele frequency ≥ 0.1 in dbSNP were shown for comparison.

Next, we examined the mutation spectrum of the mosaic variants. Similar to inherited germline variants [44] and somatic mutations reported in cancer studies [45], C>T is the most predominant type for mosaic variants (Figure 4B). We then extracted the tri-nucleotide genomic context of each variant and decomposed the matrix into mutation signatures previously identified in various types of cancers (https://cancer.sanger.ac.uk/cosmic/signatures). Single base mutation signature analysis further revealed that over 50% of the mosaic variants can be decomposed into the combination of cancer signatures 1, 5 and 30 (Figure 4C). Signatures 1 and 5 result from the age-related process of spontaneous or enzymatic deamination of 5-methylcytosine to thymine; signatures 18 and 30 result from deficient base excision repair [46]; signature 2 indicated the activation of AID/APOBEC cytidine deaminase; signatures 6 and 20 are associated with defective DNA mismatch repair; signature 22 is associated with aristolochic acid exposure; the etiology of signatures 8, 12, 19, 25 are unknown, and signatures 51 and 58 are potential sequencing artefacts. Detailed descriptions of the signatures are provided in Supplemental Text.

To explore the general relationship between the MAF of a mosaic variant and the carrier phenotype, we extracted the allele fraction and phenotypic severity information for each mosaic variant in MosaicBase. For mosaic variants in the “parent” group, we observed that the mosaic variants in parents with milder or full disease phenotypes had significantly higher MAFs than those of asymptomatic parents (P = 5.9×10^−5^ by a two-tailed Mann-Whitney U test with continuity correction, Figure 4D), which is in accordance with previous estimations [18, 20, 47]. When we considered mosaic variants in all the collected individuals, the difference became even more significant (P < 2.2×10^−16^ by a two-tailed Mann-Whitney U test, Figure 4D). These results highlighted the importance of the MAF information of mosaic variants in clinical applications such as genetic counseling.

## Discussion

MosaicBase currently contains 34,689 mosaic SNVs and indels identified in patients with noncancer diseases and their parents, as well as asymptomatic individuals, with rich information at the publication, individual, gene and variant levels. The user-friendly interface of MosaicBase allows users to access our database by multiple searching methods and the integrated genome browser.

The pathogenic contribution of mosaic variants to noncancer diseases has been increasingly recognized in the past few years. MosaicBase provides genetic and phenotypic information about 6,698 disease-related mosaic variants in 269 noncancer diseases. This database may help clinicians understand the pathogenesis and inheritance of mosaic variants and shed new light on future clinical applications, such as genetic counseling and diagnosis. On the other hand, the collection of 27,991 mosaic variants that were identified in asymptomatic individuals could be useful for understanding the genomic baseline of postzygotic mutations in the general human population. MosaicBase also integrates risk prediction from multiple computational tools for each variants. Unlike germline variants which are present in all cells of the carriers, mosaic variants are only present in a fraction of cells, in which the level of mosaic fraction can be an additional factor contributing to variant pathogenicity [18, 20]. In the future, with the increasing number of mosaic-related studies, we would expect a well-benchmarked scoring system specifically designed for predicting the deleterious probability of mosaic variants.

Of the 34689 mosaic variants collected in MosaicBase, only 0.7% to 8.7% were present in large-scale population polymorphism databases (Supp. Table 5). If we only considered common SNPs with population allele frequency (AF) higher than 0.01, the overlapping proportion further reduced to 0.1% to 0.7%. This suggested that MosaicBase provided a unique set of human genetic variants which had been overlooked in previous genomic studies. Indeed, these apparently benign variants which are generated *de novo* show characteristics distinct from those of the variants that directly contribute to a disease phenotype, and also different from polymorphisms that are fixed in population under selective pressure (Figure 4). The data from MosaicBase will also encourage researchers to reanalyze existing NGS data of human diseases by mosaic variant calling tools, such as MosaicHunter [48], Mutect2 [49], and Strelka [50], to identify previously ignored disease causative variants.

In the future, our team will update MosaicBase regularly by collecting and reviewing new publications in PubMed and publications submitted through our online submission system. After each update, we will update the statistics and release update reports on the website. We plan to further improve the user interface of MosaicBase and add new analysis tools based on feedback from the community.

## Authors’ contributions

AYH, LW, and XY, conceived the idea of building the database about mosaic variants, XZ, XY, and CY designed and implemented the website. XY, CY, LX, YT, YD, QW, and JL collected the data. MW, and AYY assisted in the website development. XY and XZ analyzed the data from the website. XY, CY, XZ, and AYH wrote the manuscript. AYH and LW led the project.

## Competing interests

The authors declare no competing interests.

## Acknowledgments

This work was supported by the National Natural Science Foundation of China (No. 31530092) and the Ministry of Science and Technology of China (2015AA020108). We thank Dr. Lei Kong, Dr. Ge Gao, and Mr. Dechang Yang from the Centre for Bioinformatics, Peking University, for their efforts in helping to set up the hardware environments and maintaining the MosaicBase server. We thank Drs. Sijin Cheng and Jinpu Jin for their suggestions for the database.

## Supplemental Materials

### Supplemental Text

Literature curation and variant collection.

Detail description of single base substitution signatures.

Web Resources.

### Supplemental Tables

Supp. Table S1: Field description for the table of publication information.

Supp. Table S2: Field description for the table of individual information.

Supp. Table S3: Field description for the table of variation information.

Supp. Table S4: Summary for mosaic SNVs and indels in noncancer diseases and asymptomatic individuals in MosaicBase.

Supp. Table S5: Comparisons between postzygotic mosaic variants and human genetic variations identified by large-scale sequencing projects.

**Supplemental References**

## Notes

http://49.4.21.8:8000/

http://mosaicbase.cbi.pku.edu.cn/

