## supplement materials for "MosaicBase: A Knowledgebase of Postzygotic Mosaic Variants in Noncancer Diseases and Asymptomatic Human Individuals"


Author(s) Names

Xiaoxu Yang[1,#,a], Changhong Yang[2,3,4#,b], Xianing Zheng[4,#,c], Luoxing Xiong[5,d], Yutian Tao[4,6,e], Meng Wang[1,f], Adam Yongxin Ye[1,5,g], Qixi Wu[7,h], Yanmei Dou[1,i], Junyu Luo[4,j], Liping Wei[1,*,k], August Yue Huang[1,*,l]

[1]Center for Bioinformatics, State Key Laboratory of Protein and Plant Gene Research, School of Life Sciences, Peking University, Beijing 100871, China

[2]Department of Bioinformatics, Chongqing Medical University, Chongqing, China

[3]College of Life Sciences, Beijing Normal University, Beijing 100875, China

[4]National Institute of Biological Sciences, Beijing 102206, China

[5]Peking-Tsinghua Center for Life Sciences (CLS), Academy for Advanced Interdisciplinary Studies, Peking University, Beijing 100871, China

[6]Chinese Academy of Medical Sciences and Peking Union Medical College, Beijing 100730, China

[7]School of Life Sciences, Peking University, Beijing 100871, China

### Xiaoxu Yang, Changhong Yang, and Xianing Zheng contributed equally to this work

* (Wei L) and (Huang A Y)


#### Running title:

*Yang X et al / Knowledgebase for Mosaic Variants*

### Supplemental Material Contents

#### Supplemental Text

Literature curation and variant collection.
Detail description of single base substitution signatures.
Web Resources.

#### Supplemental Tables

Supp. Table S1: Field description for the table of publication information.
Supp. Table S2: Field description for the table of individual information.
Supp. Table S3: Field description for the table of variation information.
Supp. Table S4: Summary for mosaic SNVs and indels in noncancer diseases and asymptomatic individuals in MosaicBase.
Supp. Table S5: Comparisons between postzygotic mosaic variants and human genetic variations identified by large-scale sequencing projects.

#### Supplemental References

### Supplemental Text

**Literature curation and variant collection**

The query string for PubMed was "((mosaic[Title/Abstract] OR mosaicism[Title/Abstract] OR neurogenesis[Title/Abstract] OR (post zygotic)[Title/Abstract] OR somatic[Title/Abstract] ) AND ((next generation sequencing) OR (deep sequencing) OR (sequencing) OR mutational) NOT cancer[Title] NOT tumor[Title/Abstract] NOT tumour[Title/Abstract] NOT plant NOT leukaemia NOT *oma NOT virus NOT transgen* NOT knockout NOT knockin NOT (carcinoma) NOT (sarcoma) NOT review[Publication Type]) AND ("1989/01/01"[Date - Publication] : "2018/06/01"[Date - Publication])", and a total of 5919 results were returned. We further excluded all publications about cancer-related mosaic mutations or studies on non-human organisms by manual check the title and abstract. For the remaining publications, we scrutinized the main text as well as supplemental information to further confirm their relevance to our study. As a result, 383 journal research articles passed all the filters. We manually collected detailed information at publication-, individual-, and variant-level. Genomic coordinates of each variants were computed from their cDNA accession using Mutalyzer [1] or directly obtained from the publication, and further converted between hg19/RCh37 and hg38/GRCh38 version via UCSC liftover.

**Detail description of single base substitution signatures**

S1: An endogenous mutational process initiated by spontaneous or enzymatic deamination of 5-methylcytosine to thymine which generates G:T mismatches in double stranded DNA. Failure to detect and remove these mismatches prior to DNA replication results in fixation of the T substitution for C. S1 is clock-like in that the number of mutations in most cancers and normal cells correlates with the age of the individual. Rates of acquisition of S1 mutations over time differ markedly between different cancer types and different normal cell types. These differences correlate with estimated rates of stem cell division in different tissues and S1 may therefore be a cell division/mitotic clock.

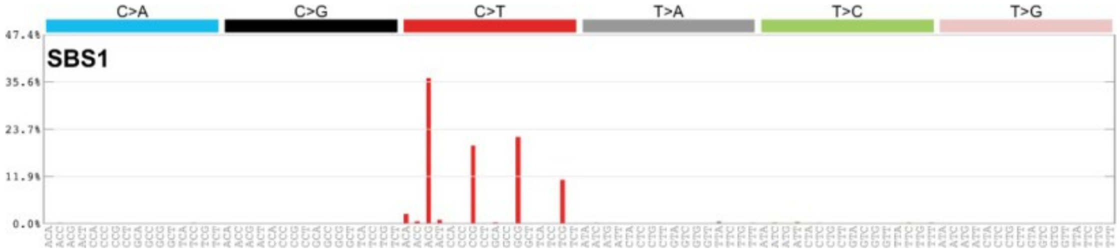


S2: S2 is usually found in the same samples as S13. It has been proposed that activation of AID/APOBEC cytidine deaminases in cancer may be due to previous viral infection, retrotransposon jumping, or tissue inflammation.

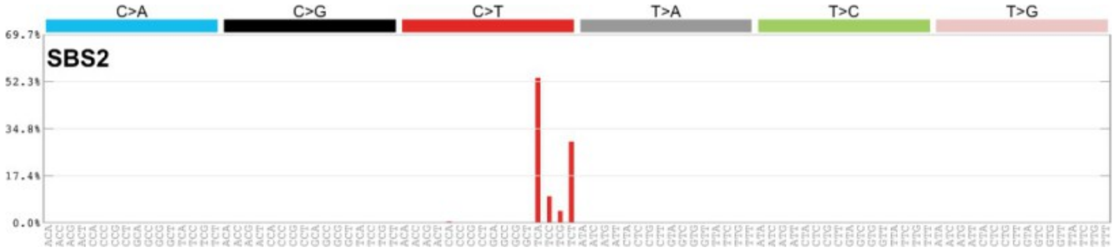

S5: S5 is clock-like in that the number of mutations in most cancers and normal cells correlates with the age of the individual. Rates of acquisition of S5 mutations over time differ between different cancer types and different normal cell types. These differences do not clearly correlate with estimated rates of stem cell division in different tissues nor with differences in S1 mutation rates. S5 may be contaminated by S16.

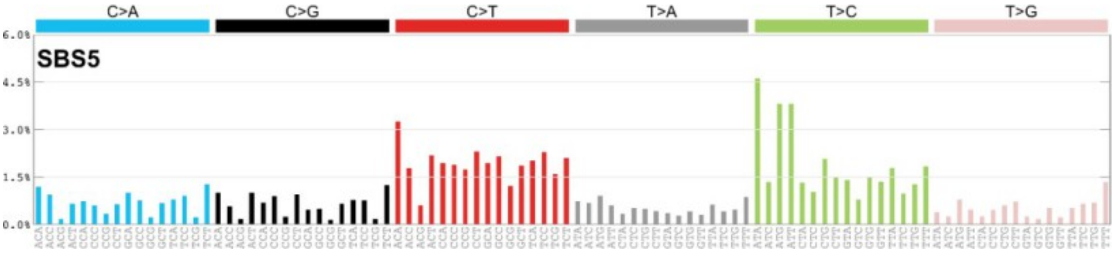


S6: S6 is one of seven mutational signatures associated with defective DNA mismatch repair (with microsatellite instability, MSI) and is often found in the same samples as other MSI associated signatures: S14, S15, S20, S21, S26, and S44.

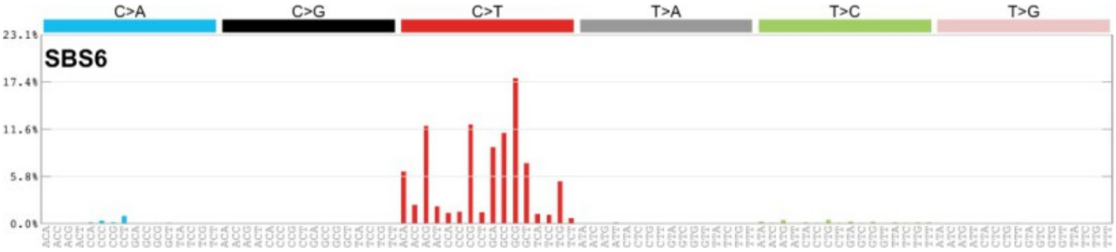


S8: The etiology of S8 is unknown, it is associated with CC>AA mutations.

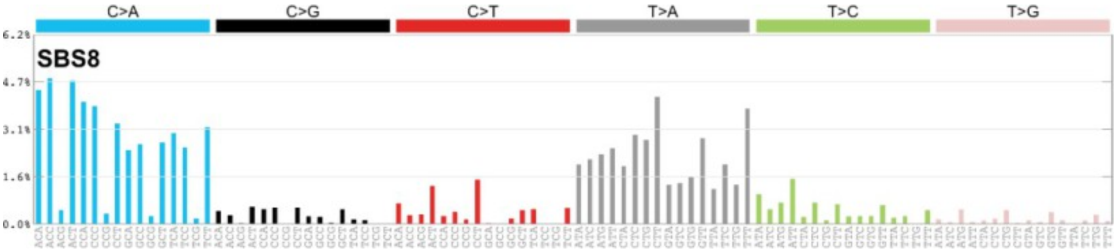


S12: The etiology of S12 is unknown.

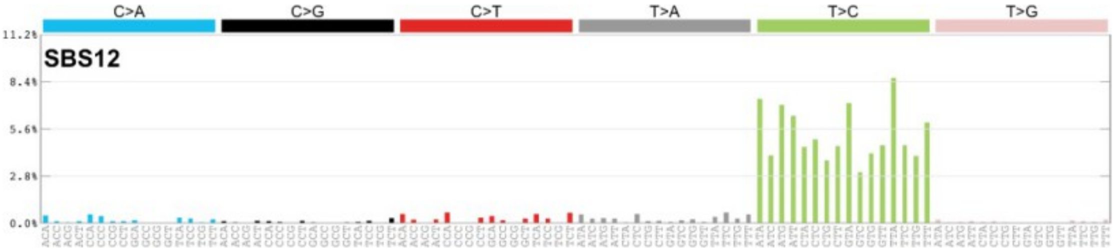


S18: S18 is similar in profile to S36 which is associated with defective base excision repair.

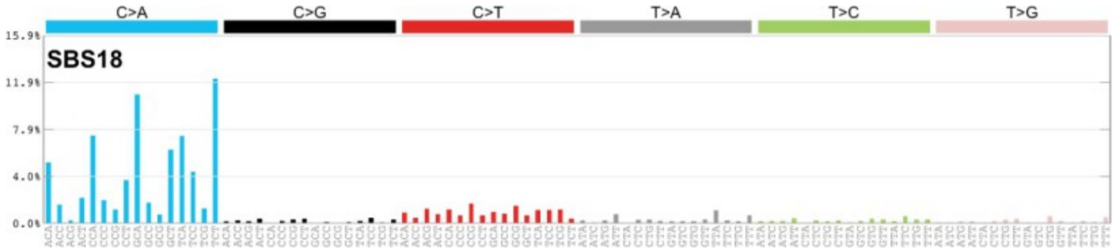


S19: The etiology of S19 is unknown.

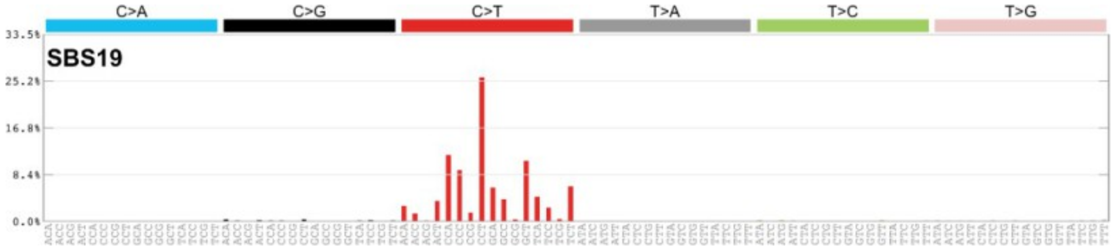


S20: S20 is one of seven mutational signatures associated with defective DNA mismatch repair (MSI) and is often found in the same samples as other MSI associated signatures: S6, S14, S15, S21, S26, and S44.

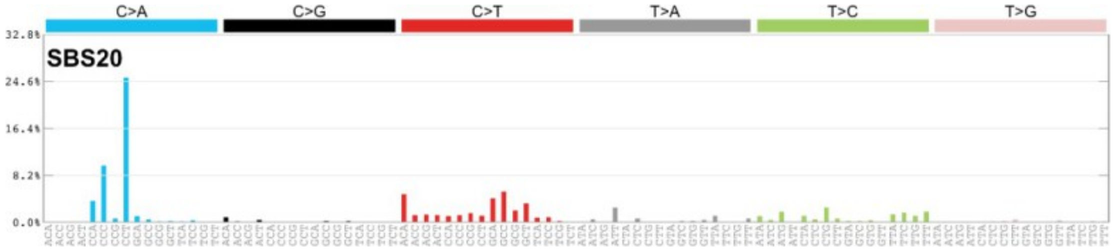


S22: S22 has been found in experimental systems exposed to aristolochic acid.

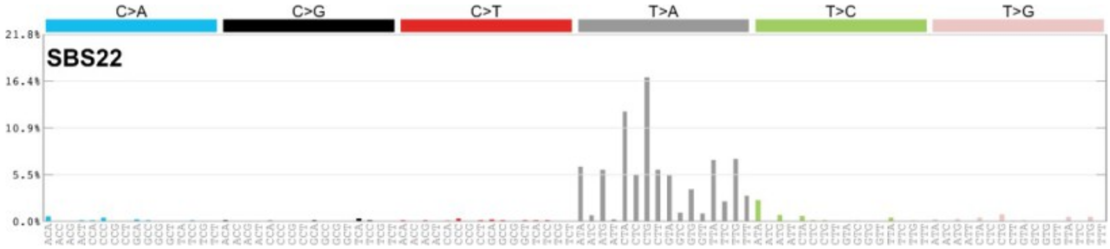


S25: The etiology of S25 is unknown. This signature was identified in Hodgkin's cell lines.

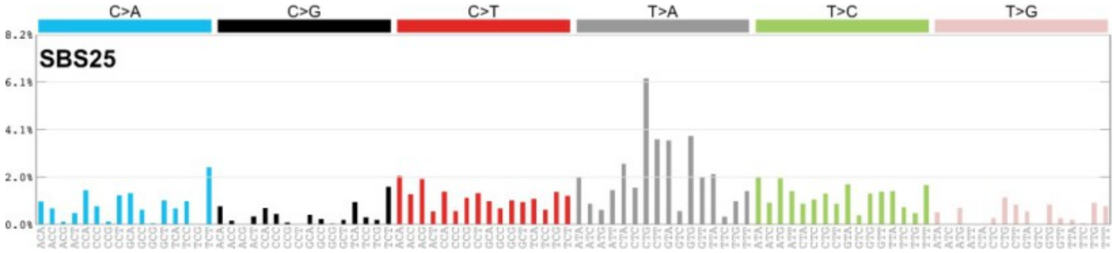


S30: S30 is due to deficiency in base excision repair.

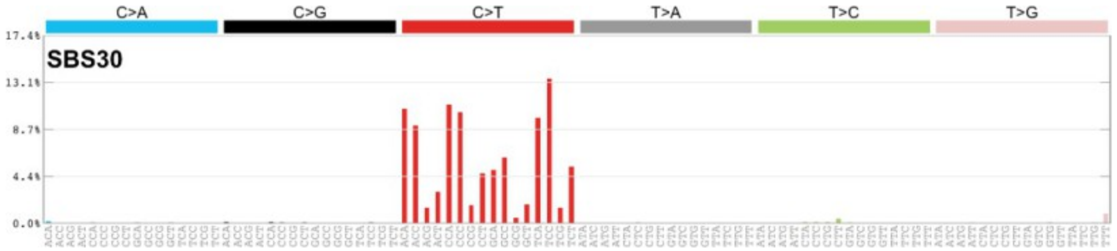


S51: S51 is a potential sequencing artefact.

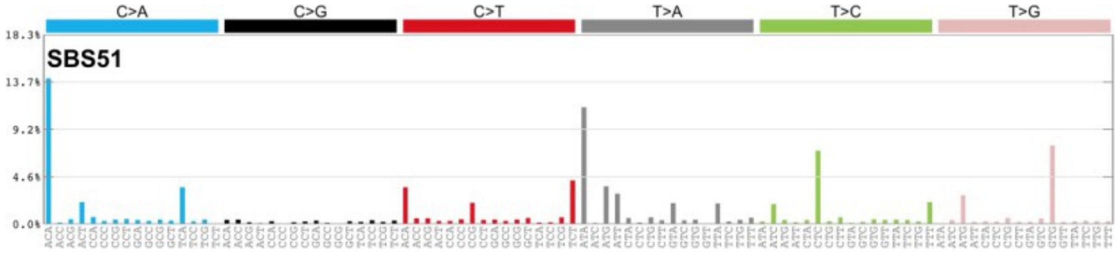


S58: S58 is a potential sequencing artefact.

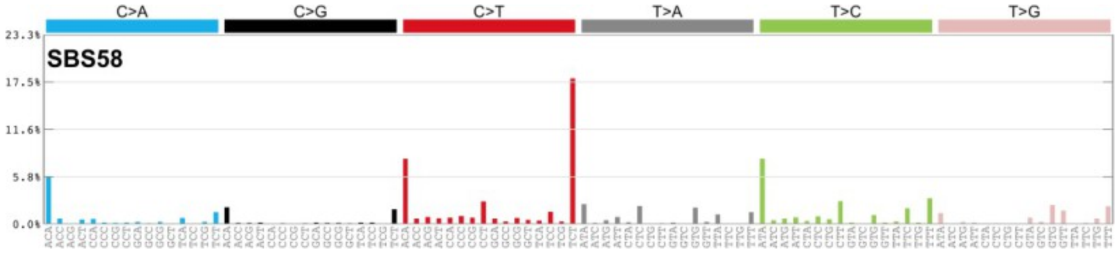


**Web Resources**

MosaicBase, http://mosaicbase.cbi.pku.edu.cn/

Biodalliance, http://www.biodalliance.org/

Disease Ontology, http://disease-ontology.org

Mutalisk, http://mutalisk.org/

Mutalyzer, https://mutalyzer.nl/
